# Subsistence of sib altruism in different mating systems and Haldane’s arithmetic

**DOI:** 10.1101/2022.06.16.496366

**Authors:** József Garay, Villő Csiszár, Tamás F. Móri

## Abstract

The moral rule “Risk your life to save your family members” is, at the same time, a biological phenomenon. The prominent population geneticist, J.B.S. Haldane told his friends that he would risk his life to save two drowning brothers, but not one – so the story goes. In biological terms, Haldane’s arithmetic claims that sib altruism is evolutionarily rational, whenever by “self-sacrifice” an altruistic gene “rescues”, on average, more than one copy of itself in its lineage. Here, we derive conditions for evolutionary stability of sib altruism, using population genetic models for three mating systems (monogamy, promiscuity and polygyny) with linear and non-linear group effect on the siblings’ survival rate.

We show that for all considered selection situations, the condition of evolutionary stability is equivalent to Haldane’s arithmetic. The condition for evolutionary stability is formulated in terms of genetic relatedness and the group effect on the survival probability, similarly to the classical Hamilton’s rule. We can set up a “scale of mating systems”, since in pairwise interactions the chance of evolutionary stability of sib altruism decreases in this order: monogamy, polygyny and promiscuity.

Practice of marrying and siblings’ solidarity are moral rules in a secular world and in various religious traditions. These moral rules are not evolutionarily independent, in the sense that the subsistence of sib altruism is more likely in a monogamous population.

**Highlights:**

- Haldane’s arithmetic is introduced
- Conditions for evolutionary stability of sib altruism are given
- Evolutionary stability is equivalent to Haldane’s arithmetic in the studied model
- Generalized Hamilton’s rules are formulated

## 1. Introduction

Darwin (1871) already raised the possibility that moral has an evolutionary origin (Boehm 2012; Brooke 2013; Garay et.al, 2014, 2019; Wilson 2010; Ridley 1997). Altruism is a key issue in human morality. For instance, the partial version of golden rule (Skyrms 1996): “*Risk your life to save your family members if you want them to save your life*” (Garay et. al. 2019) can be considered as an altruistic biological behaviour (Sussman & Cloninger 2011) and as a moral rule^1^ at the same time.

There are three families of evolutionary models used to study this problem: kin selection, group selection and population genetic models. Nowadays, kin selection and group selection theorists form two circles (Nowak et al. 2010; Abbot et al. 2011; Okasha 2010; Birch & Okasha 2015; De Vladar & Szathmáry 2017; Van Veelen et al. 2017), based on their different fundamental views. The kin selectionist circle focuses on the genetic relatedness between recipient and actor (Haldane 1955; Hamilton 1964), while the group selectionist circle focuses on the advantages of group living (Boehm 2012; Simon 2014). We note that these theories are not necessarily antagonistic, since if all interactions are additive or pairwise, then both theories yield the same predictions (Ven Veellen 2009; 2020). Otherwise, however, the relationship between these two theories is problematic (e.g. Ven Veellen 2009; 2020). This is one reason why we think that more detailed, so necessarily particular models are needed to shed new light on the evolutionary stability of altruism in diploid sexual populations.

Here we use a population genetic model that can account for the frequency-dependent interactions between genotypes within the families, and the genetic background of the inheritance of the altruistic phenotype (Cavalli-Sforza & Feldman 1978, Feldman & Cavalli-Sforza 1981; Maynard Smith 1980; Spencer & Feldman 2005; Uyenoyama & Feldman 1980; Uyenoyama et al. 1981). Recently, we introduced a concept of evolutionary stability in a general population genetics model in which the relative frequencies of diploid genotypes are the states of the model. Since the survival game between juvenile excludes that the genotype distribution is at Hardy–Weinberg equilibrium in the next parental generation, our model is based on the mating table (Garay et al. 2019). Our population genetics model incorporates two of the main motives of the kin and group selection theories: a) the genetic relatedness between siblings and phenotypic composition of the family are determined by the mating and the inheritance systems; b) the survival probability of a sibling stuck in a death trap depends on the phenotypic composition and the size of the group – now the family. We adapt the definition of evolutionary stability by Maynard-Smith and Price (1973): A strategy is evolutionarily stable, if a rare enough mutant genotype cannot invade the monomorphic resident population. We note that our population genetic model describes the genotype changes from generation to generation, and genotype exclusively determines the phenotype, thus the phenotypic view of the verbal evolutionary stability concept can be used directly. In our previous paper, we considered only monogamy, and all interactions were additive. There we found that classical Hamilton’s rule implies evolutionary stability of altruism. In this paper we consider other mating systems and non-additive group effect on the siblings’ survival.

During the investigation of our concrete selection situations by a population genetic model, we come across the following two well-known ideas of evolutionary theory: Hamilton’s rule and Haldane’s idea on kin selection (see Section 2). The classical Hamilton’s rule states that social behaviours will evolve whenever *rb* > *c*, where *r* denotes relatedness between social partners, *b* the effect of the behaviour on the recipient of the behaviour, and *c* the effect of the behaviour on the actor of the behaviour (Hamilton 1964). This classical Hamilton’s rule can be generalized on the basis of Price’s equation (Price 1970; Gardner et al., 2011), but for different selection sutations the precise mathematical formulas of Hamilton’s rule are different (see Table 1, Gardner et al. 2011). Moreover, Van Veelen (2005) points out that the results obtained by the Price equation are sometimes not correct (Van Veelen et al. 2010, 2012). This is the another reason why we think that mathematically precise models are needed. So we follow van Veelen and his coathurs’s (2010) call by treating population genetic models with rigour.

**Table 1.** Probabilities of the progeny genotypes in different types of matings, calculates by Mendelian rules.

| Mating | Progeny $G_1$ [$a, a$] | Progeny $G_2$ [$a, A$] | Progeny $G_3$ [$A, A$] |
| --- | --- | --- | --- |
| $G_1 \times G_1$ | 1 | 0 | 0 |
| $G_1 \times G_2$ | $\frac{1}{2}$ | $\frac{1}{2}$ | 0 |
| $G_1 \times G_3$ | 0 | 1 | 0 |
| $G_2 \times G_2$ | $\frac{1}{4}$ | $\frac{1}{2}$ | $\frac{1}{4}$ |
| $G_2 \times G_3$ | 0 | $\frac{1}{2}$ | $\frac{1}{2}$ |
| $G_3 \times G_3$ | 0 | 0 | 1 |

## 2. Introducing Haldane’s arithmetic

The question of the evolutionary rationale of this moral rule is a part of the history of population genetics. Legend^2^ has it that in the nineteen-fifties, Haldane told his friends in a pub one evening that he would risk his life to save two brothers, but not one, and similarly, he would risk his life to save eight cousins, but not seven (Haldane 1955; McElreath & Boyd 2007). Haldane made a distinction by his genetic relatedness to his relatives. Genetic relatedness is the average proportion of shared genes, i.e. the probability that a gene picked randomly from each individual at the same locus is identical by descent. Under monogamy, the genetic relatedness between full sibs and cousins is 1/2 and 1/8, respectively. In other words, a gene of Haldane is identical to one parental gene of his two brothers and to one grandparental gene of his eight cousins, on average. Surprisingly, although Haldane’s idea is well-known and widely cited, as far as we know it has not been formalized and investigated. In this paper we propose a reformulation of Haldane’s idea in terms of gene balance: In anthropomorphic wording, *Haldane’s arithmetic* claims: *The relative frequency of the altruistic gene increases, if by “self-sacrifice”, it has “rescued” on average more than one copy of itself in its lineage*.

Before we come to the main point of our paper, we show that Haldane’s arithmetic implies the classical Hamilton’s rule (but not conversely, see Appendix 1). Let us suppose the closest relatives of Haldane get in a death trap. Haldane can try to save them one by one. Let the success probability of every attempt be equal to *p* (benefit of recipient), and suppose that every rescue attempt can cause the death of the rescuer with probability *q* (cost of actor). Then the average number of rescued relatives during his life is *p*/*q* (see Appendix 1). Denote by *r* the genetic relatedness between all the rescued people and Haldane. Thus, during its life, one altruistic gene has rescued *rp*/*q* of its copy by self-sacrifice. We can assume that the non-altruistic gene surely survives, thus Haldane’s arithmetic leads to the classical Hamilton’s rule, i.e. *rp*/*q* > 1.

Note that Haldane’s arithmetic does not reckon with the possibility that should the altruistic individual also get in a death trap, his altruistic siblings try to help him too. Neither does it calculate with the survival probability of an individual in a death trap without altruistic help. We also remark that Haldane’s arithmetic is general in the sense that the probabilities *p* and *q* may be arbitrary, e.g., they can depend on the number of altruistic family members, consequently, on the size of the groups as well. One of the main differences between Haldane’s and Hamilton’s view is that Haldane concentrated on his own lineage. However, both are in essence a cost/benefit condition, only Haldane’s arithmetic does not take directly into account the genetic relatedness.

## 3. A population genetic model

Here we concentrate on two aspects of sib altruism. Firstly, genetic relatedness between siblings depends on the mating system and on the level of promiscuity in the population. In human societies, in the vast majority of cases, polygamy is accepted (Gray 1998; Starkweather & Hames 2012) and the typical rate of non-paternity is estimated about 10%, depending on the ethnographic conditions, but with a wide range from 0.4% to 50% (Anderson 2006). For the sake of simplicity, and because of the insufficiency of our information on the mating system and on the level of promiscuity during human evolution, here we concentrate only on three extreme mating systems: monogamy without promiscuity, promiscuity (Starkweather & Hames 2012) (where the female randomly chooses whom to mate with, consequently the offspring of a promiscuous female are all half sibs) and polygyny without promiscuity (Gray 1998) (a male has more than one female partners at the same time, who only mate with him). Secondly, filial altruism belongs to group selection, since interactions happen only in a genetically determined group, a family. For instance, the survival rate under predator attacks is higher in families with more altruistic defenders (Garay et al. 2014). Our population genetics model incorporates two of the main motives of these theories: a) the genetic relatedness between siblings and phenotypic composition of the family are determined by the mating and the inheritance systems (Garay et. al. 2019); b) the survival probability of a sibling stuck in a death trap depends on the phenotypic composition and the size of the group – now the family.

### Population genetic model

We study the evolutionary stability of sib altruism in the framework of a population genetic model (Garay et. al. 2019). We consider a diploid, sexual population with equal sex ratio (½ – ½), with no sexual selection, i.e., females and males differ only in their sex, there is no mating preference. A dominant-recessive autosomal allele pair determines the phenotype of each individual. For the sake of simplicity we only consider the case when the resident allele *a* is recessive under the mutant *A* allele, i.e. the resident genotype [*a, a*] is altruistic and the genotypes [*a, A*] (primary mutant) and [*A, A*] (secondary mutant) are non-altruistic. In other words, a mutation immediately appears in the phenotype. We emphasize that in the resident sexual population each individual has self-sacrificing phenotype coded by a totally homozygote genotype.

In the considered large population the genotype distribution among the offspring follows the Mendelian rules (see mating Table 1), thus secondary mutants can only be born from mutant-mutant intercourses. The corresponding genotype frequencies in the population are *y*_1_, *y*_2_, *y*_3_, resp. These frequencies are the same for females and males. We suppose that the mutants are sufficiently rare, thus we consider the case where *y*_1_ → 1. Suppose *y*_1_ = 1 − *ε*, then *y*_2_ ∼ *ε*, and *y*_3_ = *O*(*ε*^2^).

We concentrate on self-sacrificing sib altruism, when altruistic sibs help their siblings, while non-altruistic siblings do not. The altruistic interaction takes place only within the families (which are of size *n*, in addition to the parents), and decreases and increases the survival rate of the donor and the recipient, respectively. An altruistic sib’s survival probability decreases by *q* = *c*(*n* − 1), where *c* is a positive constant. If somebody has exactly *k* altruistic siblings, her/his survival probability increases to

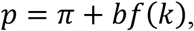

where *π* denotes the base survival rate (before the act of altruism), *b* is a positive constant and the function *f* describes the group effect on the survival probability. Regarding this group effect, we consider two situations.

a. *Pairwise interactions*. If the success of rescue depends on each sibling’s willingness to help, but does not depend on the composition of the family, we get the additive model setting

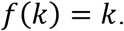
b. *Group dependent survival rate*. In this case, the success of rescue depends on the number of altruistic siblings in the family. Now we consider a function *f*(*k*) similar to that used by Hauert and coauthors (Hauert et al. 2006)

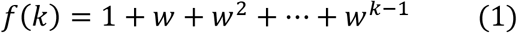

with a positive parameter *w*. The parameter *w* determines the group effect on the survival rate, namely, when *w* > 1, *w* = 1, and *w* < 1, our function *f* describes a synergetic, additive, and discounted group effect, respectively. In the synergetic case the overall effect of the group is larger than the sum of individually provided benefits, while in the discounted case the reverse inequality is valid. We assume that benefits from different sources as well as costs accumulate. Note that all offspring of suitable type are involved in sibling-sibling interactions, not only those who survive. We remark that Hauert et al. (2006) divided *f*(*k*) by the family size *n* as well, and their model differs from ours at several further points: in our model the altruist does not provide a benefit to herself, and the amount of per capita beneficial payoff is fixed, rather than inversely proportional to the number of the offspring. Moreover, in our model the cost is proportional to the number of beneficiaries rather than being fixed.

Our objective is to find sufficient conditions in terms of *b* and *c* for the resident genotype to be evolutionary stable. This concept is defined as follows. Let *Y*_1_, *Y*_2_, *Y*_3_ denote the average number of resident, primary, and secondary mutant individuals, resp., in the next generation. The resident genotype is called *evolutionary stable*, if mutation cannot spread provided it is sufficiently rare (Maynard Smith and Price, 1973); that is, there exists a threshold *ε* > 0 such that *Y*_1_ > *y*_1_(*Y*_1_ + *Y*_2_ + *Y*_3_) whenever *y*_1_ ≥ 1 − *ε*. In other words, the growth rate of the resident is higher than the average growth rate in the population. So, we are exclusively interested in whether the altruistic or the non-altruistic variant of a randomly chosen parental gene will result in more surviving copies on average, provided the resident population is homozygous altruist. Note that the definition does not change if *Y*_1_, *Y*_2_, *Y*_3_ denote the average number of type [*a, a*], [*a, A*], and [*A, A*] individuals, resp., in a family chosen at random. In our computation we will use *Y*_1_, *Y*_2_, *Y*_3_ in this latter sense. Since *Y*_3_ is negligible compared with *Y*_1_ and *Y*_2_, evolutionary stability of the resident is implied by the inequality (1 − *y*_1_)*Y*_1_ > *y*_1_*Y*_2_.^3^

### Conditions for evolutionary stability in different mating systems

We start with the derivation of the conditions for the evolutionary stability of altruism. (For mathematical details see Appendix 2). We focus on the three mating systems mentioned above.

### Monogamy

The condition of evolutionary stability reads as

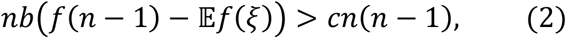

where *ξ* denotes the random number of altruistic siblings of an individual in a family with parents *G*_1_ × *G*_2_. It has a binomial distribution with parameters *n* − 1 and 1/2. We note that *f*(*n* − 1) − 𝔼*f*(*ξ*) measures the group effect.

In the additive model, in accordance with Hamilton’s rule and (Garay et. al. 2019), we obtain that sib altruism is evolutionarily stable if

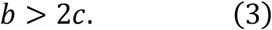

In a discounted or synergetic model with *w* ≠ 1 and fixed family size *n* we get the condition

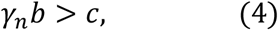

where

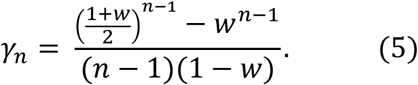

The parameter region satisfying condition (4) is visualized in Figures 1a and 1b for family sizes *n* = 4 and *n* = 8, respectively. For mathematical details see Appendix 2.A.

**Figure 1.**
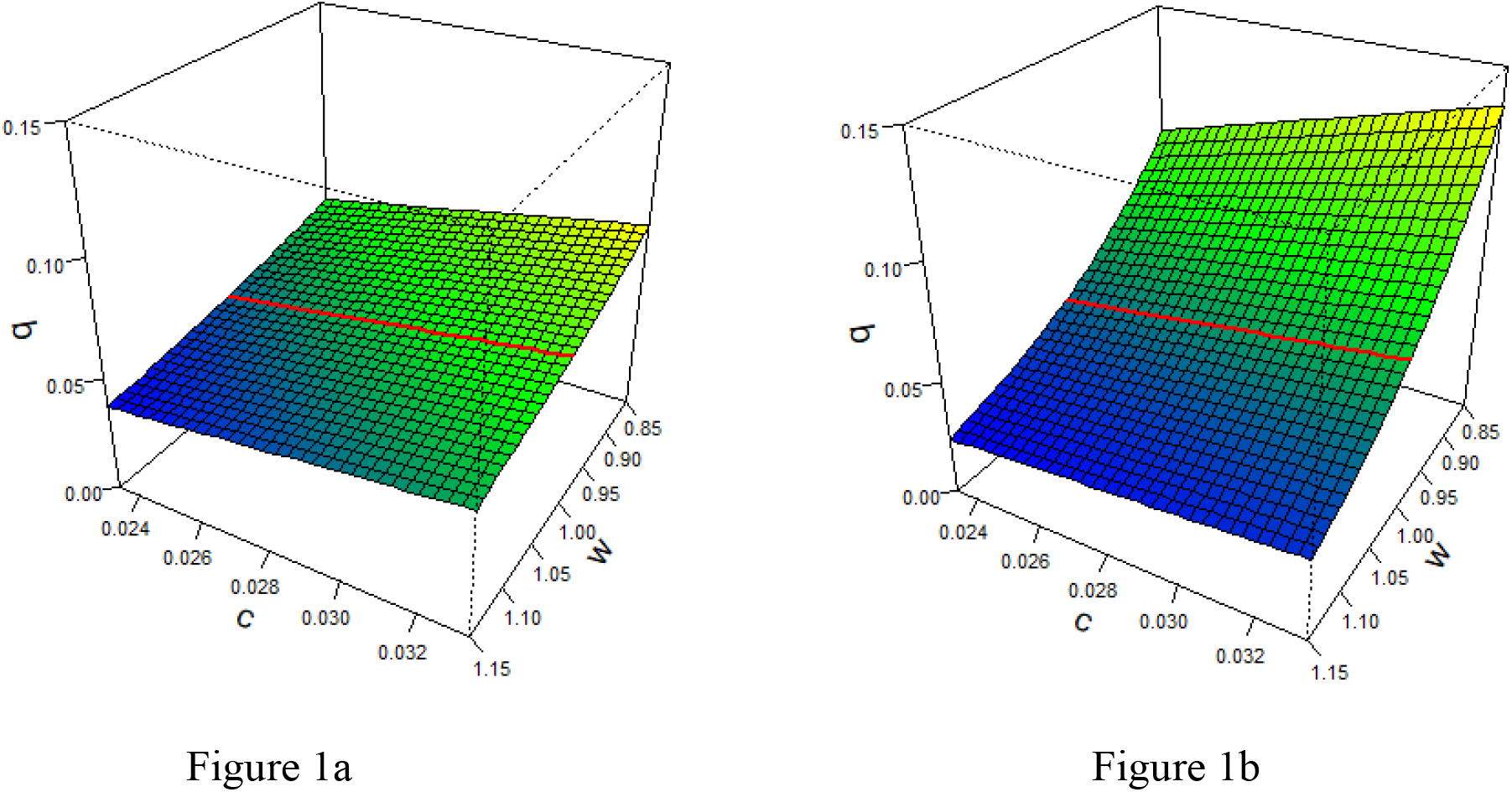
When does altruism appear remunerative: Monogamy. In Figures 1a and 1b the threshold value for *b* is plotted against *c* and *w*, where the family sizes are *n* = 4 in Figure 1a, and *n* = 8 in Figure 1b (for the details of the model see the bottom of the box). Every point in the cubes corresponds a triplet (*w, c, b*). Points above the colored surface indicate models where altruism is evolutionarily stable, or equivalently, Haldane’s arithmetic also predicts evolutionarily stability of altruism. The red line corresponds to the additive case *w* = 1, where Hamilton’s rule holds. That is, Hamilton’s rule implies the evolutionary stability of sib altruism in pairwise interactions. The red lines are located in the same positions of the two cubes, because in the additive case the condition for the evolutionary stability of sib altruism does not depend on the family size. On the other hand, in the discounted and synergetic cases (*w* ≠ 1), the family size does have a palpable effect on the evolutionary stability of altruism, in accordance with the group selection theory. Comparing Figures 1a and 1b we can see that in the synergetic case (*w* > 1) a bigger family size allows the evolutionary stability of sib altruism to occur with lower benefit *b*, while in the discounted case (*w* > 1) the effect of the family size is just the opposite. This is heuristically obvious, because cost is linearly proportional to family size. The same figures apply to the case of **total promiscuity** with the only modification that the points of the vertical axis represent *b*/2 instead of *b*. Thus, the basic idea of kin selection is in effect, since the average genetic relatedness between siblings in polygamous families is just the half of that in monogamous families (see also Figure 3). *Model*: The family size (offspring size) is equal to *n*. An altruistic sibling increases the survival probability of each of her *n* − 1 siblings while her own survival probability decreases by (*n* − 1)*c*. If an individual has *k* altruistic siblings, then the overall increment of her survival probability is *b*(1 + *w* + *w*^2^ + … + *w*^*k*−1^), where the positive parameter *w* controls the group effect: it is discounted for *w* < 1, synergetic for *w* > 1, while *w* = 1 is the purely additive case. Ranges of parameters in both figures are: 0.85 ≤ *w* ≤ 1.15, 0.0233 ≤ *c* ≤ 0.0333, 0 ≤ *b* ≤ 0.15.

### Total promiscuity

The condition of evolutionary stability now reads as

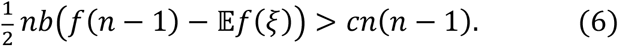

In the additive model, this is simply

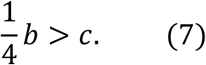

Observe that this condition is just the classical Hamilton’s rule.

In a discounted/synergetic model with fixed family size *n* and benefit function (1) the condition is

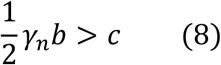

where *γ*_*n*_ is defined in (5). Figure 1 applies to this case as well. For mathematical details see Appendix 2.B.

### Polygyny without promiscuity

In this setup, families consist of one male and *k* females living together. Each female is assumed to produce *n* offspring. Thus each sib has full and half siblings, too. Moreover, each sib has perfect information whether his/her sib in danger is a full or a half sibling. Thus the altruistic interaction can depend on the genetic relatedness of the interacting siblings. If an individual gets support from *ℓ* altruistic full siblings and *m* half-sibs, then her/his survival probability increases by *g*(*ℓ, m*). Plausible candidates for the function *g* can be obtained as follows. In the monogamous case, a benefit function *f* can be interpreted by thinking of altruistic siblings acting one by one, with the *i*th one increasing the survival probability by *b*Δ*f*(*i*), where Δ*f*(*i*) = *f*(*i*) − *f*(*i* − 1) ≥ 0. In the present case, suppose that somebody has *ℓ* altruistic full siblings and *m* altruistic half-sibs. The *i*th altruistic sibling increases the survival probability of the beneficiary by *b*_1_Δ*f*(*i*) or *b*_2_Δ*f*(*i*) depending on whether she is a full sibling or a half-sib. If the altruistic siblings act in a random order (with equal probabilities of all possible rosters), the average benefit is

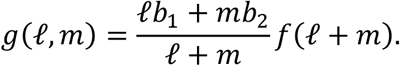

We will use the benefit function *f* defined in (1). The particular case of *f*(*k*) = *k*, that is, *g*(*ℓ, m*) = *ℓb*_1_ + *mb*_2_, will again be referred to as *additive*. We also suppose that the cost is *c*_1_ or *c*_2_ depending on whether the interaction takes place between full or half siblings, respectively. In this mating system our evolutionary stability condition looks rather complicated. The condition reads as

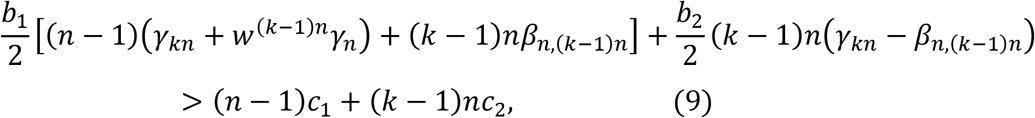

where *γ*_*n*_ and *γ*_*kn*_ are defined in (5), and *β*_*n*,(*k*−1)*n*_ is defined as

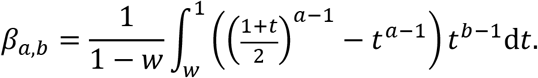

We also consider the particular case when there is no difference between the attitude toward full siblings or half-sibs, that is, *b*_1_ = *b*_2_ = *b* and *c*_1_ = *c*_2_ = *c*. Then condition (9) for the evolutionary stability of altruism reads as

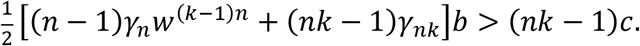

In the additive model the condition of evolutionary stability is reduced to the following inequality.

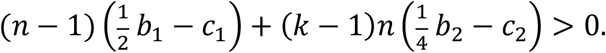

This condition is in accordance with the inclusive fitness theory of Hamilton (1964): multipliers 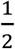 and 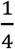 are just the coefficients of genetic relationship for full siblings and half-sibs, resp., and the multipliers *n* − 1 and (*k* − 1)*n* are proportional to the expected number of full sibling and half-sib pairs, resp. Furthermore, if there is no difference between the attitude towards full siblings and half-sibs, that is, *b*_1_ = *b*_2_ = *b*, and *c*_1_ = *c*_2_ = *c*, then the above condition is simplified to

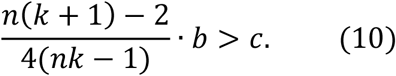

We emphasize that condition (10) coincides with the classical Hamilton’s rule. Some parameters for which condition (9) holds are visualized in Figure 2. For mathematical details see Appendix 2.C.

**Figure 2:**
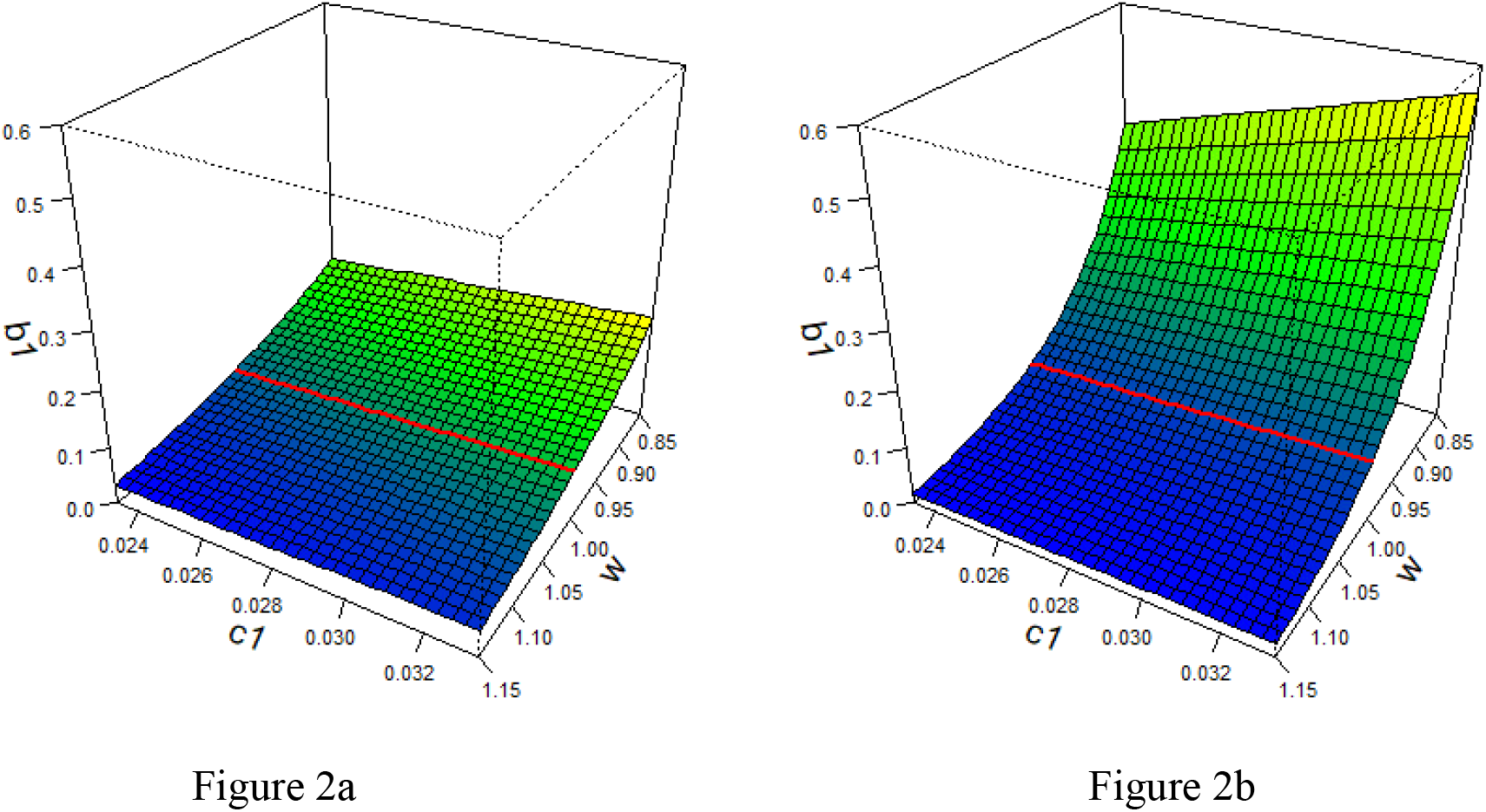
When does altruism appear remunerative: Polygyny without promiscuity. Here we consider families where there are one male and *k* females living together and each female gives birth to *n* children. In Figures 2a and 2b the threshold value for *b*_1_ is plotted against *c*_1_ and *w*, while *k, n*, and the ratios *b*_2_/*b*_1_ and *c*_2_/*c*_1_ remain fixed (for the explanation of the parameters see the bottom of the box). The family parameters are: *n* = 4, *k* = 2 in Figure 2a, and *n* = 4, *k* = 4 in Figure 2b. Every point in the cubes corresponds a triplet (*w, c*_1_, *b*_1_). Points above the colored surface indicate models where altruism is evolutionarily stable or, equivalently, where Haldane’s arithmetic predicts evolutionary stability of altruism. The red line corresponds to the additive case (*w* = 1), where Hamilton’s rule holds. That is, Hamilton’s rule implies the evolutionary stability of sib altruism in pairwise interaction. It is not hard to compute (see expression (8) in Appendix 2.C) that the average genetic relatedness among the 8 siblings in the family of Figure 2a is 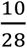, and the same quantity is equal to 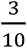 for the 16 siblings in the family of Figure 2b. The figures show that the effect of genetic relatedness can suppress the effect of the group living advantage based on the family size, except the case where a relatively strong synergy is present (the colored surface lies higher in Figure 2b than in Figure 2a, at least for values of *w* less than or close to 1). *Model*: An altruistic sibling increases the survival probability of each of her *n* − 1 full siblings and *n*(*k* − 1) half-sibs, while its own survival probability decreases by (*n* − 1)*c*_1_ + *n*(*k* − 1)*c*_2_. If an individual gets support from *ℓ* altruistic full siblings and *m* half-sibs, then her survival probability increases by 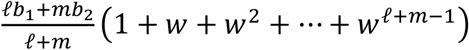, where the positive parameter *w* controls the group effect: it is discounted for *w* < 1, synergetic for *w* > 1, and *w* = 1 is the purely additive case. Ranges of parameters in both figures are: 0.85 ≤ *w* ≤ 1.15 and 0.0233 ≤ *c*_1_ ≤ 0.0333 (the same as in Figures 1a–b), 0 ≤ *b*_1_ ≤ 0.6. Moreover, *b*_2_ = 0.8*b*_1_ and *c*_2_ = 0.8*c*_1_ are fixed.

For the sake of simplicity, here we considered the case of equal and non-random family size. For generalizations of the results to random family sizes, see Appendix 2.

### Comparison of different mating systems

Now we are in the position to ask which mating system favors mostly the subsistence of sib altruism? Figures 3a and 3b suggest that – except in the case where a relatively strong synergy is present – the chance of evolutionary stability of sib altruism decreases (i.e. the parameter domain where sib altruism is evolutionarily stable shrinks) in this order: monogamy, polygyny and promiscuity.

**Figure 3:**
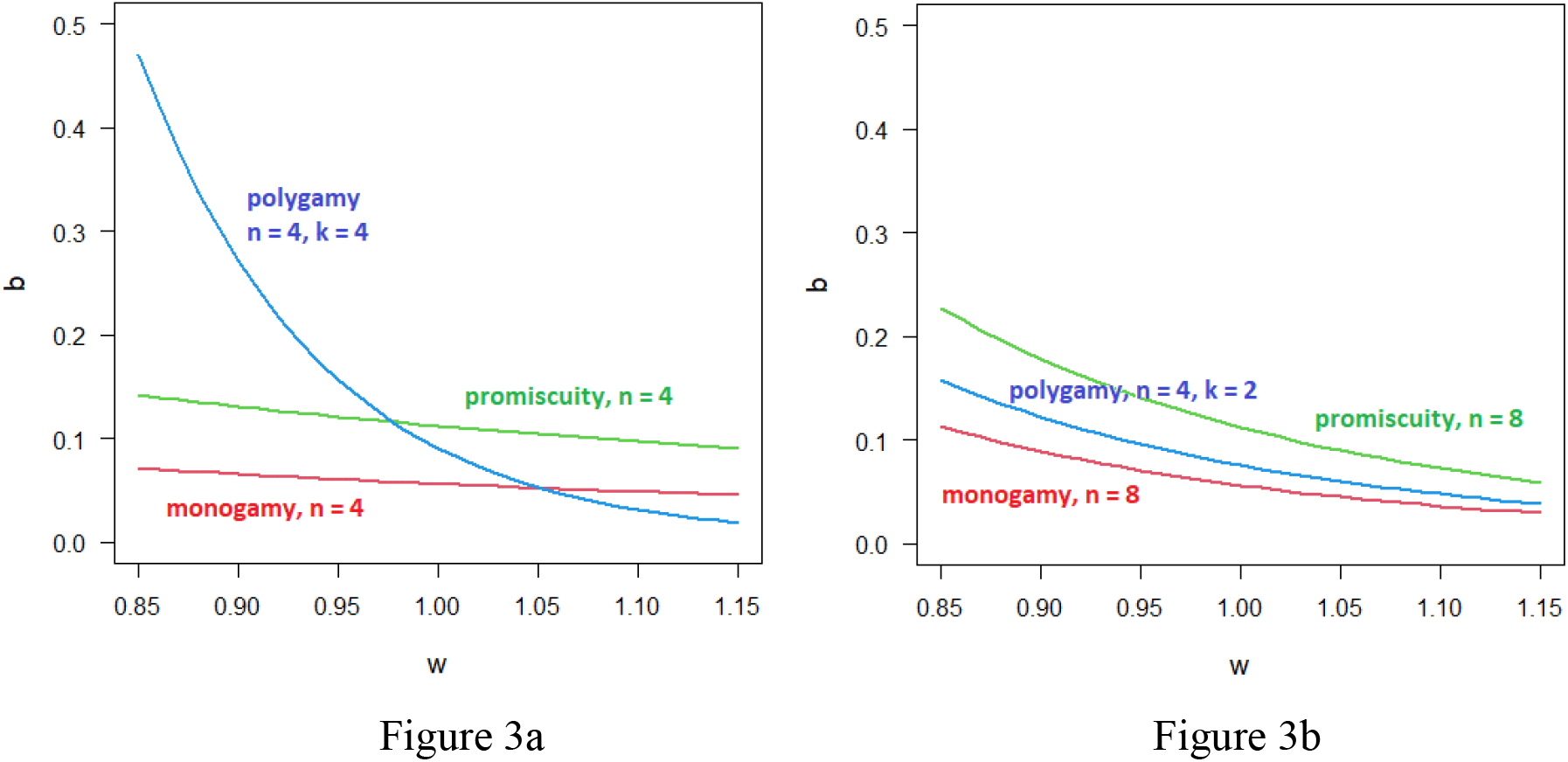
Effects of average genetic relatedness and group living. Figure 3a and 3b demonstrate the effects of average genetic relatedness and those of the group living advantage. In Figure 3a each female has 4 offspring independently of the mating system. As we have already mentioned in Figure 1, sib altruism can be evolutionarily stable at a smaller benefit *b* under monogamy than under promiscuity. However, when the group living advantage is highly synergetic (in our example *w* > 1.05) polygamy can outperform monogamy based on synergetic family size effect. We emphasize that when the altruistic interaction is pairwise (the benefit is additive at *w* = 1), then altruism becomes fixed more likely in monogamous than polygamous or promiscuous environment, and polygamy is more advantageous for altruism to become fixed than promiscuity. In order to exclude the effect of differences in the family size, in Figure 3b we consider three families with 8 offspring. In all three cases the considered families are phenotypically equivalent, thus no asexual model could detect any difference between the mating systems. On the other hand, in our population genetic model an efficiency order of mating systems appears even at equal family sizes, since monogamy, polygyny and promiscuity contribute to the chance of evolutionary stability of sib altruism in decreasing order. In the figures the condition for *b* is plotted against the group effect parameter *w* in the cases of monogamy (red curve) and total promiscuity (green curve), resp. The cost parameter *c* is set to 0.028. In the case of polygyny (polygamy) without promiscuity (blue curve), *b*_1_ is plotted as a function of *w*, and *b*_2_ = 0.8*b*_1_, *c*_1_ = 0.028, *c*_2_ = 0.8*c*_1_ are assumed. For the models see Figures 1 and 2.

### Haldane’s arithmetic is equivalent to the condition of evolutionary stability

Having derived the evolutionarily stability conditions, we show in Appendix 3 that these conditions are equivalent to Haldane’s arithmetic in all considered selection situations. More precisely, we derive that in all mating systems in question, if the interaction is pairwise (i.e. additive), then the classical Hamilton’s rule is a special case of Haldane’s arithmetic, which is in turn equivalent to the evolutionary stability condition. However, when costs and benefits are not additive, then it is Haldane’s arithmetic (but not the classical Hamilton’s rule) that is equivalent to the evolutionary stability of altruism.

### Generalized form of Hamilton’s rule

As we have already mentioned in the Introduction, the classical Hamilton’s rule can be generalized on the basis of Price’s equation (Gardner et al., 2011). Although Gardner and his coaouthors offered a different method for the calculation of the actual version of Hamilton’s rule, here we already have evolutionary stability conditions, which we now reformulate in a way to resemble the form of the classical Hamilton’s rule (see Appendix 4). Namely, *rB* > *C*, where *r* is the genetic relatedness, *B* = *B*(*r*) is the synergetic group effect modified average benefit of the recipient and *C* is the average cost to the donor^4^. Under monogamy and total promiscuity the genetic relatedness between siblings is uniform, i.e. ½ and ¼, respectively. Observe that our conditions (4) and (8) can be read as

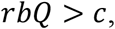

where

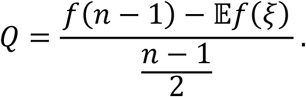

Thus, our generalized Hamilton’s rule takes account of the synergetic group effect with the multiplier *Q* and this effect is different from the parameters of the classical Hamilton’s rule. Under polygyny without promiscuity, each sib has full and half siblings, too. Formally, our condition of the evolutionary stability of altruism (9) reads as

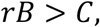

where

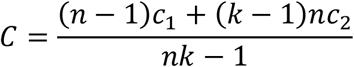

is the average cost; the average genetic relatedness within the families is

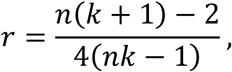

and *B* is the synergetic group effect combined with benefit parameters. Here the discounted/synergetic group effect cannot be separated from the parameters of the classical Hamilton’s rule. In the additive model where the group does not take effect, we have

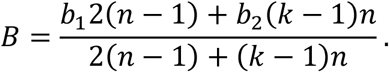

For the details of the derivation, see Appendix 4.

In summary, by applying the notion of evolutionary stability in population genetics we provide a new method for the derivation of the actual form of the generalized Hamilton’s rule. Observe that the detailed population genetics model takes account of the basic motives of both kin and group selection theories (i.e., the genetic relatedness and the synergetic group effect). We think that the importance of the formulation of Hamilton’s rule lies in getting insight into the effect of the genetic relatedness. Finally, we note that the above versions of Hamilton’s rule are equivalent to Haldane’s arithmetic, and both represent a cost/benefit condition for the evolutionary stability of altruism.

## 5. Discussion

In this paper, we use one of the basic tenets of biology, i.e., that evolution is genetically determined, so we can assume that individuals’ behaviour is fully geneticly determined. Although this study was motivated by a human moral rule (de Waal 2006), it strictly belongs to the theory of evolutionary population genetics. Our main observation is that genetic relatedness (as claimed by kin selection theory) and group effect in inetarction (as claimed by group selection theory) together determine the formal conditions of evolutionary stability of altruism. Our results call the attention to the possibility that under the umbrella of Haldane’s arithmetic, the harmony of kin and group selection theories can be approached, at least in a mechanistic population genetic model.

From the point of view of kin selection theory, monogamy ensures the highest genetic relatedness between siblings (e.g. Davies et al. 2016, Cornwallis et al. 2010). For instance, Lukas and Clutton-Brock (2012) found that the evolution of cooperative breeding in mammals has been restricted to socially monogamous species, which currently represent 5 percent of all mammalian species. Moreover, it seems that the lifetime monogamy was the universal ancestral state for obligate eusociality (Boomsma 2009, 2013). Thus our population genetics results are in harmony with the biological facts and kin slection theory, as well.

We emphasize that two of the main phenomena of group selection theory are used in the population genetic model presented hereby. The first one is group formation, which, in our case, is defined by the genetic system (i.e., the mating system and inheritance rules) determining the number of altruistic siblings in the families. However, the group formation process affects the condition of the evolutionary stability of altruism by changing the genetic relatedness within the families according to the different mating systems. The second one is the non-linear group effect on the survival probability, given by the functions *f* and *g*. However, our present research question does not take account of other phenomena of group selection theory, namely, the competition between families (e.g. Birch and Okasha, 2015) and group-level phenomena, e.g., competition between groups formed by unrelated families, moreover fission, fusion, and extinction of groups of families (e.g., Garay et al., 2014, or Simon, 2014). Clearly, considering these phenomena in a new pure population genetics model is a task of future works, and the following question is to be answered: are Haldane’s arithmetic and the condition of evolutionary stability equivalent in those models? We think this problem analytically hard when multiple loci determine the altruism.

Haldane’s arithmetic, similarly to Hamilton’s rule, is a cost/benefit condition for the evolutionary stability of altruism, but it does not directly take genetic relatedness into account. The clear difference between Haldane’s view and the “inclusive fitness” view is the following. Haldane’s arithmetic concentrates on a parental gene, and the genetic relatedness among its offspring determines the number of copies of the parental gene in question. In other words, from the genes’ point of view, genetic relatedness is an important factor in the future but it is not regarding the past. In anthropomorphic wording: The selfish gene “is only interested” in the survival rate of its own copies (lineage), and “does not care about” the survival rate of other (perhaps identical) genes.

## 6. Conclusions

### A contemporary example for human sib altruism and the possible statistical investigations

Is there a way to check on real data that altruistic help is more likely between full siblings than between half-sibs? In our previous paper (Garay et al. 2019) we have already suggested that “self-sacrificing” could be observed in kidney transplantations, when the live donor is a family member (Guttman et al. 2016). The estimated rate of transplants relying on live donations from family members was 37% in Taiwan, 99% in Japan, and 66% in South Korea (Tanaka et al., 2004), 80% in Mexico (Crowley-Matoka, 2016). However, in Egypt about 70–90% of kidney transplants used kidney from Cairo’s thriving black market, so the donation may also depend on the ethnographic conditions (Hamdy, 2012). In the US, between 1987 and 2012, related living donors were parents 81.7%, full siblings 6.9%, half siblings 0.8%, identical twins 0.1%, and other relatives 10.5% (Van Arendonk et al., 2013). Between 1987 and 2017, 35% were biologically unrelated, 8% were half-siblings or other biological relatives, 13% were parents, 16% were offspring, 29% were full siblings, and 0.2% were identical twins (Muzaale et al., 2020). However, these data cannot be used directly to study the question above, mainly for the following reasons: Firstly, we should take into account how many full and half siblings a patient had. Secondly, the donor candidates may be turned down due to ABO/HLA incompatibilities with the recipient.

For studying the above question the following data are surely needed about each recipient: How many living relatives, full siblings and half-sibs she/he has? Among them who are willing to be a donor? How many of the offers are eligible? If kidney was transplanted, from what donor: relative, non-relative, or even non-living (e.g., kidney from an accident)? With a recipient and donor follow-up one can attempt to estimate the survival rates. A possible study would face several difficulties due to hidden variables. On the one hand, the composition of the recipient’s family is sociologically determined, and sociological aspects may affect the decisions of possible donors. On the other hand, compositions of families differ in different racial and ethnicity groups. For instance, in the US, the percent of all children under 18 who live with at least one half sibling, is the least among Asian (3%), and the highest among black (14%) in 2004 (Kreider and Fields, 2010). However, the number of full siblings are approximately equal in different races (see Muzaale et al., 2020, Table 1). Statistical inference must be carried out with extra caution as the presence of relevant hidden variables may lead to Simpson type paradoxes (Wagner, 1982).

### Evolutionay stability in sexual and asexual models

The problem whether the result of evolution is the same in sexual and asexual populations goes back to Maynard Smith (1981), who raised the following question: “*Will a sexual population evolve to an ESS?*” (The classical ESS corresponds to asexual populations.) The answer is positive if the interaction is given by a well-mixed matrix game between genotypes, and in a sexual population, in a dominant-recessive inheritance system, *n* alleles at one autosomal locus determine *n* possible pure individual phenotypes, and each pure phenotype is obtained as the phenotype of a homozygote (Garay and Varga, 2003). However, the answer is negative if the interaction is not well-mixed between genotypes (e.g., if it only happens in the family), and in a sexual population there is a dominant-recessive inheritance system (Garay et al., 2019). Based on those, in selection situations where the interaction is not well-mixed in a sexual population (see the group selection models), the population genetic model should give deeper insight. For instance, from the perspective of asexual models, our selection situations of monogamy and polyandry with total promiscuity are the same (since the group sizes are equal), thus an asexual model cannot make difference between them. Consequently, our results point out that the predictions of an asexual model cannot be directly applied to sexual populations.

### Moral rules are not necessarily evolutionarily independent

The practice of marrying with a certain level of promiscuity and the solidarity between siblings are moral rules in a secular world and in various religious traditions. Evolution theory highlights that these moral rules are not independent, since in monogamous groups of our ancestors, the subsistence of sib altruism was more likely than in polygynous or polygamous groups. Finally, the present paper was motivated by a human moral rule among family members, where we assumed that behaviour was genetically determined. However, human culture has an effect on human behaviour, too. In the theory of evolution, this problem can be studied by cultural evolution models (e.g., Creanza et al., 2017). Thus, the following (open) question arises: can Haldane’s arithmetic play a role in cultural evolution models?

## Software used

Figures were drawn in R version 4.0.2.

## Declarations

### Ethics approval and consent to participate

Not applicable.

### Consent for publication

Not applicable.

### Availability of data and materials

Not applicable.

### Authors’ contributions

JG and TFM designed the study. VCs and TFM analyzed the model, VCs made the figures and JG, VCs and TFM wrote the article.

## Acknowledgements

We thank Andy Gardner, Ádám Kun, István Scheuring, András Szilágyi, Tibor Standovár and Zoltán Varga for their valuable comments on the earlier version of this paper.

**Authors’ information (optional)**

## Appendix 1. Formalization of Haldane’s arithmetic

Suppose some of your relatives get in a death trap. You can try to save them one by one. The success probability of every attempt is *p*, and the successive attempts are independent. On the other hand, every rescue attempt can cause the death of the rescuer with probability *q*. We are looking for conditions on *p* and *q* such that the average number of rescued people is at least *R*.

Suppose the rescuer dies after *Y* attempts (even the last, fatal rescue attempt can be successful). This random variable has geometric distribution with expectation 1/*q*. Conditionally on *Y*, the distribution of the number of saved people is binomial with parameters *Y* and *p*. Hence, by the law of total expectation we have that the average number of rescued people is *p*𝔼(*Y*) = *p*/*q*, and it is greater than or equal to *R* if and only if *p* ≥ *Rq*.

Let us carry out a cost-benefit analysis from the point of view of a gene. Suppose there are *n* people to save. A person can get in trouble several times, thus *n* can be arbitrarily large, even infinity. Denote by *r*_*i*_ the genetic relatedness between the altruistic individual and the *i*th person to rescue. (Note that the degree of genetic relatedness is defined as the probability that a pair of randomly sampled homologous alleles are identical by descent.) Suppose that the rescue attempts keep going on until a successful trial or the death of the altruistic individual. Then the probability that the first relative is rescued at the *k*-th attempt is

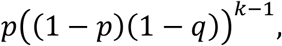

thus the overall probability of successful rescue is

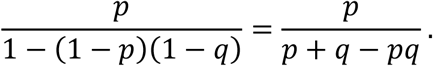

This quantity has to be multiplied by 1 − *q* for the probability that the altruistic individual also survives. Turning to the *i*th relative to save, the probability that she/he will be rescued is

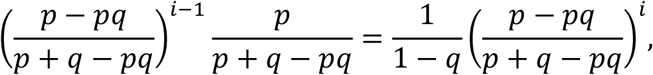

thus the average number of rescued copies of the gene under consideration is

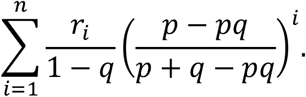

Moreover, the probability that the altruistic individual will finally survive is

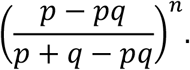

Hence the cost-benefit balance is

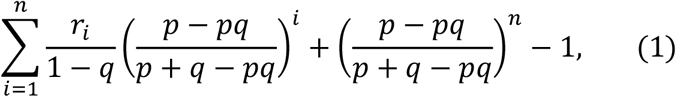

and Haldane would willingly die for his relatives if (1) is strictly positive, that is, if

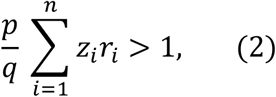

where

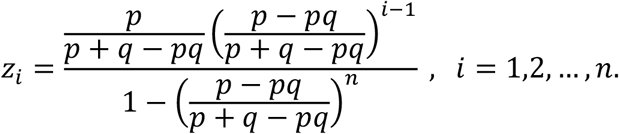

(If *n* = ∞, the denominator becomes 1, thus it can be omitted.) This is what we call *Haldane’s arithmetic*. Note that the sum in the left-hand side of (2) is a weighted average of the degrees of genetic relatedness, as the weights *z*_*i*_ add up to 1.

Particularly, when the order of relatives waiting for rescue is uniform at random, the formulae above are still valid as conditional (on the order) probabilities and expectations. Let 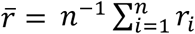, the arithmetic average of genetic relatedness, then (2) simplifies to 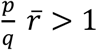 1 by the law of total expectation, hence we get Hamilton’s rule as a special case.

## Appendix 2. Evolutionary stability in population genetic model

Here we focus on monogamy, promiscuity, and polygyny.

### Appendix 2.A. Monogamy

The case of monogamous families has already been analyzed in detail for additive benefit (that is, *f*(*k*) = *k*, see Garay et al. (2019)) and Hamilton’s rule was found to imply the evolutionary stability of sib altruism.

In every family *n* children are born, where *n* can be random, but supposed to be independent and identically distributed random variables for different families. In a randomly selected family the number of genotype [*a, a*] offspring is equal to *n* with probability 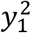 (resident– resident type family); all other possibilities are of smaller order probability. Hence the conditional expectation of the number of surviving offspring is

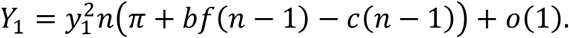

Primary mutant children can be born in families where one parent is resident or primary mutant type and the other one is an arbitrary mutant. The overwhelming majority of such families are of resident–primary mutant type. In those families there are no children of secondary mutant type. Every child is resident or primary mutant with equal probability, and the number *ξ* of altruistic siblings whose support would increase the child’s survival probability has (conditionally on *n*) binomial distribution with parameters *n* − 1 and 1/2. Thus

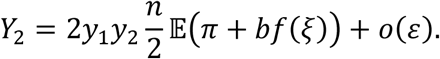

Altruism is evolutionary stable if (1 − *y*_1_)*Y*_1_ > *y*_1_*Y*_2_, that is, *y*_2_*Y*_1_ > *y*_1_*Y*_2_, because 1 − *y*_1_ ∼ *y*_2_. This yields

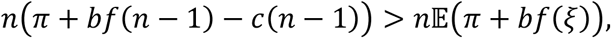

or, equivalently,

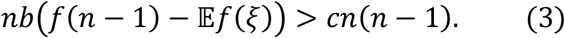

If *n* is random, we have to take expectation in both sides of (3).

In the *additive* model (that is, *f*(*k*) = *k*), 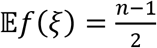, thus we obtain that sib altruism is evolutionarily stable if *b* > 2*c*, in accordance with Hamilton’s rule.

In a *discounted* or *synergetic* model, with

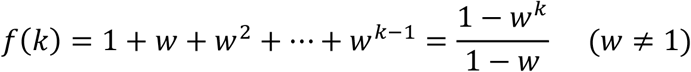

and fixed family size *n* we get

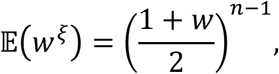

thus

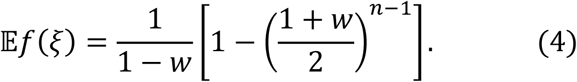

hence the condition is *γ*_*n*_*b* > *c*, where

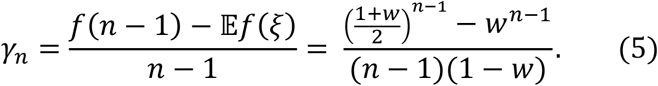

This quantity shows how much a non-altruistic mutant decreases the average group effect *per capita*. Since

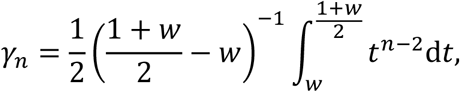

it tends decreasingly to zero as *n* → ∞ from 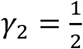, provided the model is discounted, i.e., *w* < 1. In the presence of synergy, i.e., *w* > 1, *γ*_*n*_ → ∞ increasingly as *n* → ∞. If *n* is fixed, *γ*_*n*_ is an increasing function of *w*.

### Appendix 2.B. Total promiscuity

Suppose females live in total promiscuity. Every female produces *n* children, each from randomly and independently chosen fathers. Progeny size *n* can be random; in that case they are independent and identically distributed random variables for different females. The population is considered so large that the *n* fathers can be supposed all different; thus siblings are half-sibs.

Consider a random mother. The types of her children are independent and identically distributed (but this distribution depends on the type of the mother). Firstly, let us compute the average number of her [*a, a*] type offspring up to a remainder of *o*(1). The principal term in *Y*_1_ comes from the case where both parents are of resident type. In the case of random offspring sizes let us consider the model conditionally, with offspring size supposed known. The (conditional on *n*) expectation of the number of surviving offspring is

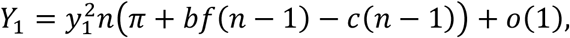

like in the monogamy case, since the probability that a given mother meets at least one mutant type mate is *o*(1), negligible.

The principal term in *Y*_2_ is obtained when one of the parents is of resident type, and the other one is a primary mutant. There is a big difference between the progeny of a resident mother and a primary mutant one. Children of a resident mother can get benefit from almost all of their siblings, while only about half of the siblings of a primary mutant mother’s child are altruistic. The probability that a randomly selected child in a randomly selected family is a resident mother’s primary mutant child is 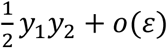, and its survival probability is *π* + *bf*(*n* − 1) + *o*(1). Though the probability of the other possibility, that is, when the selected child is a primary mutant mother’s primary mutant child, is the same (up to a remainder *o*(*ε*)), this time the survival probability is different: *π* + *b*𝔼*f*(*ξ*) + *o*(1), where *ξ* is a random variable of binomial distribution with parameters *n* − 1 and 1/2.

Altogether, we have

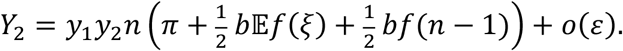

Thus, altruism is evolutionary stable if

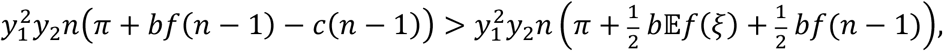

from which

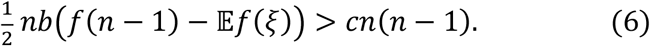

(In the case of random family size we have to take expectation in both sides.)

Note that the left-hand side of (6) is just the half of the left-hand side of inequality (3) obtained in the monogamous case, while the right-hand sides are equal. The appearance of the factor 1/2 in the left-hand side is explained by the fact that here we have almost exclusively half-sibs while in the previous case full siblings only.

Particularly, in the additive model with *f*(*k*) = *k* the condition of evolutionary stability is^5^ 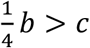, and in a discounted/synergetic model with fixed family size *n* and benefit function

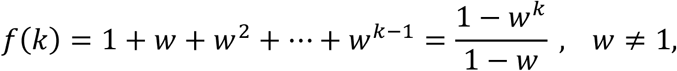

the condition is 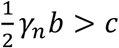, where *γ*_*n*_ is defined in (5).

### Appendix 2.C. Polygyny without promiscuity

In this model, families consist of one male and *k* females living together. The females are selected at random, so their types are independent and identically distributed according to the composition of the whole population. The females produce *n*_1_, *n*_2_, …, *n*_*k*_ offspring, respectively. Offspring sizes are independent and identically distributed random variables (including the particular case of constant offspring number, *n*_1_ = *n*_2_ = … = *n*_*k*_ = *n*). Let us allow the benefits and costs to be different between full and half-sibs: an altruistic individual increases the survival probability of each of her full siblings at the cost of *c*_1_, while this parameter is *c*_2_ in connection with half-sibs. If an individual gets support from *ℓ* altruistic full siblings and *m* half-sibs, then her survival probability increases by *g*(*ℓ, m*), where *g* is an increasing function of both arguments, with *g*(0, 0) = 0. It is plausible to suppose that *c*_1_ ≥ *c*_2_, and *g*(*ℓ* − 1, *m* + 1) ≥ *g*(*ℓ, m*), but we will not use that. Plausible candidates for the function *g* can be obtained as follows. In the monogamous case, a benefit function *f* can be interpreted by thinking of altruistic siblings acting one by one, the *i*th one increasing the survival probability by *b*Δ*f*(*i*), where Δ*f*(*i*) = *f*(*i*) − *f*(*i* − 1) ≥ 0. In the present case, suppose that somebody has *ℓ* altruistic full siblings and *m* altruistic half-sibs. The *i*-th altruistic sibling increases the survival probability of the beneficiary by *b*_1_Δ*f*(*i*) or *b*_2_Δ*f*(*i*) according that she is a full sibling or a half-sib. If the altruistic siblings act in a random order (with equal probabilities of all possible rosters), the average benefit is

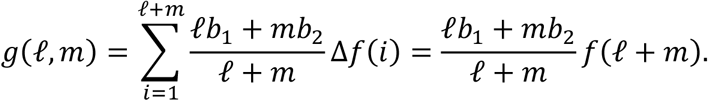

The particular case of *f*(*k*) = *k*, that is, *g*(*ℓ, m*) = *ℓb*_1_ + *mb*_2_, will be referred to as *additive*.

As before, we first make computations conditionally on the offspring sizes *n*_1_, *n*_2_, …, *n*_*k*_.

The principal term in *Y*_1_ is the contribution of families with all members being of resident type. The probability of such a family is 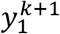, and the sum of survival probabilities in such a family is equal to

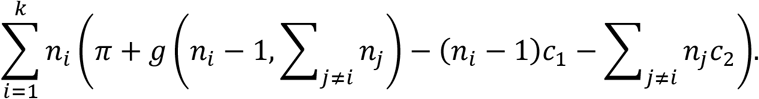

Its expectation is

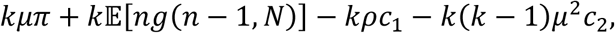

where *µ* = 𝔼*n*_1_, *ρ* = 𝔼[*n*_1_(*n*_1_ − 1)] and *n* = *n*_1_, *N* = *n*_2_ + … + *n*_*k*_.

Let us turn to genotype [*a, A*] (primary mutant). The principal term in *Y*_2_ is due to families where there is exactly one primary mutant parent, all the others are of genotype [*a, a*]. The possibility that exactly one of the females is primary mutant has probability 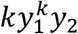 (multiplier *k* is present because any of the *k* females can be mutant). By symmetry we can suppose that this is the first female. Every offspring from the mutant female is of resident type or primary mutant with probability 1/2. A primary mutant child has exactly *N* altruistic half-sibs and a random number of altruistic full siblings. Conditionally on the offspring sizes *n*_1_, *n*_2_, …, *n*_*k*_, the expectation of the number of surviving offspring of genotype [*a, A*] is 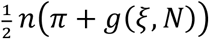 with expectation

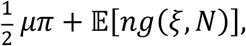

where the (conditional on the offspring sizes) distribution of *ξ* is binomial with parameters *n* − 1 and 1/2.

The other possibility is when the father is of type [*a, A*]. This occurs with probability 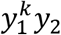. By symmetry, we may focus on his [*a, A*] type children from the first mother. Again, half of his offspring belongs to type [*a, A*] and each has (conditional) survival probability *π* + *g*(*ξ, η*), where *ξ* is the number of altruistic children of the first female, it is binomial with parameters *n* − 1 and 1/2, and *η* is the total number of altruistic offspring of the other *k* − 1 females, which is also binomial with parameters *N* and 1/2. Thus the conditional expectation of the number of surviving offspring of genotype [*a, A*] is 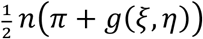 with expectation

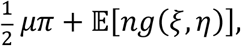

where *n, ξ* are independent of *η, N*. For the whole family this is

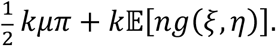

Hence the sufficient condition for altruistic behavior to be evolutionary stable is inequality

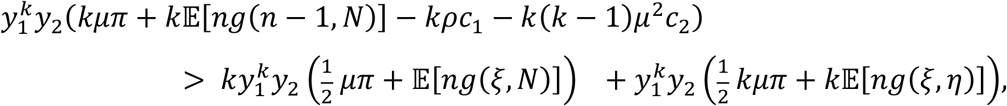

that is, after division by 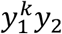 and some calculus,

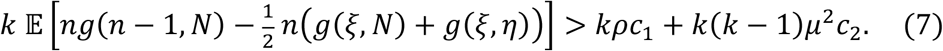

In the additive model 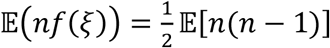 and 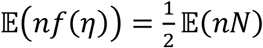, thus the left-hand side of (7) is equal to

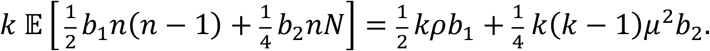

and the condition of evolutionary stability reads

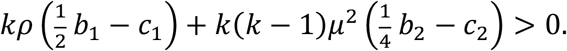

This condition is in accordance with the inclusive fitness theory of Hamilton (1964): multipliers 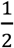 and 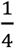 are just the coefficients of genetic relationship for full siblings and half-sibs, resp., furthermore, *kρ* and *k*(*k* − 1)*µ*^2^ are the expected numbers of full sibling and half-sib (ordered) pairs, resp.

If there is no difference between the attitude towards full siblings and half-sibs, that is, *b*_1_ = *b*_2_ = *b*, and *c*_1_ = *c*_2_ = *c*, then the above condition reads

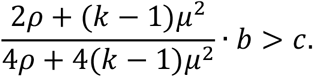

Particularly, for deterministic offspring size *n* this is

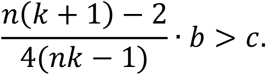

Note that the fraction on the left-hand side is just the average degree of genetic relatedness in the family. Consider an arbitrary child. She has *nk* − 1 siblings, namely, *n* − 1 full siblings and *n*(*k* − 1) half-sibs. Therefore, the probability that a randomly selected sibling carries the copy of the same parental gene is^6^

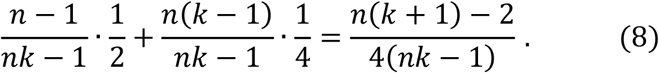

By choosing

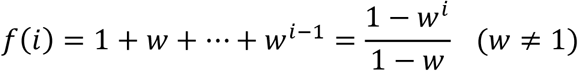

we get a discounted (*w* < 1) or a synergetic (*w* > 1) model. (For *w* = 1 we get back the additive model.) For the sake of simplicity let the offspring number *n* be constant. Then condition (7) is equivalent to the inequality

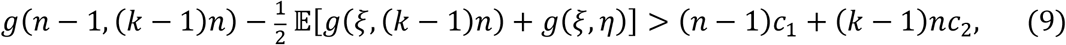

where *ξ* and *η* are independent, binomially distributed random variables with parameters (*n* − 1, 1/2) and ((*k* − 1)*n*, 1/2), resp. Here the left-hand side can be computed as follows.

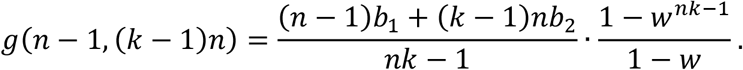

Since the distribution of *ξ* + *η* is also binomial with parameters (*nk* − 1, 1/2), and the conditional distribution of *ξ* given *ξ* + *η* = *m* is hypergeometric with parameters (*nk* − 1, *n* − 1, *m*), we get

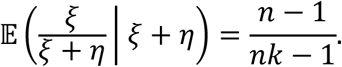

Now by (4) we have

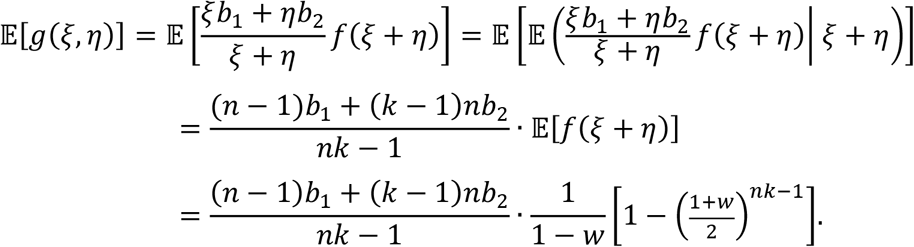

Finally, for 𝔼[*g*(*ξ*, (*k* − 1)*n*)] such a simple expression does not appear to exist.

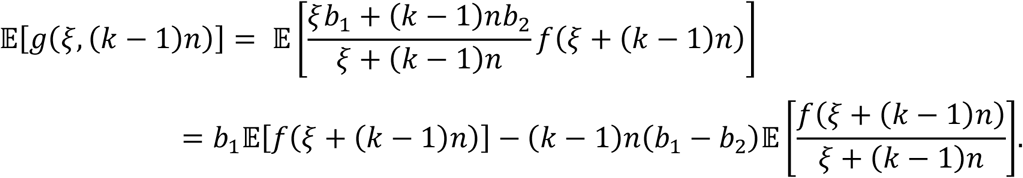

Here

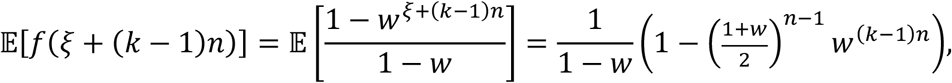

and

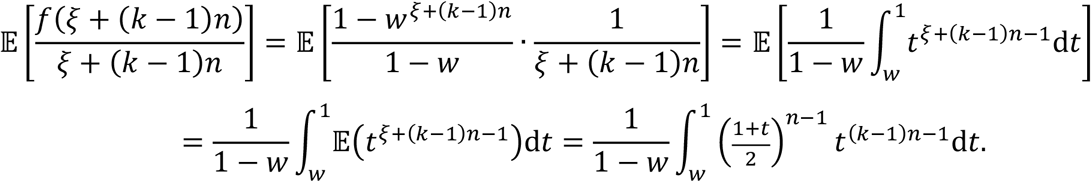

By all these, in the left-hand side of (9) we get

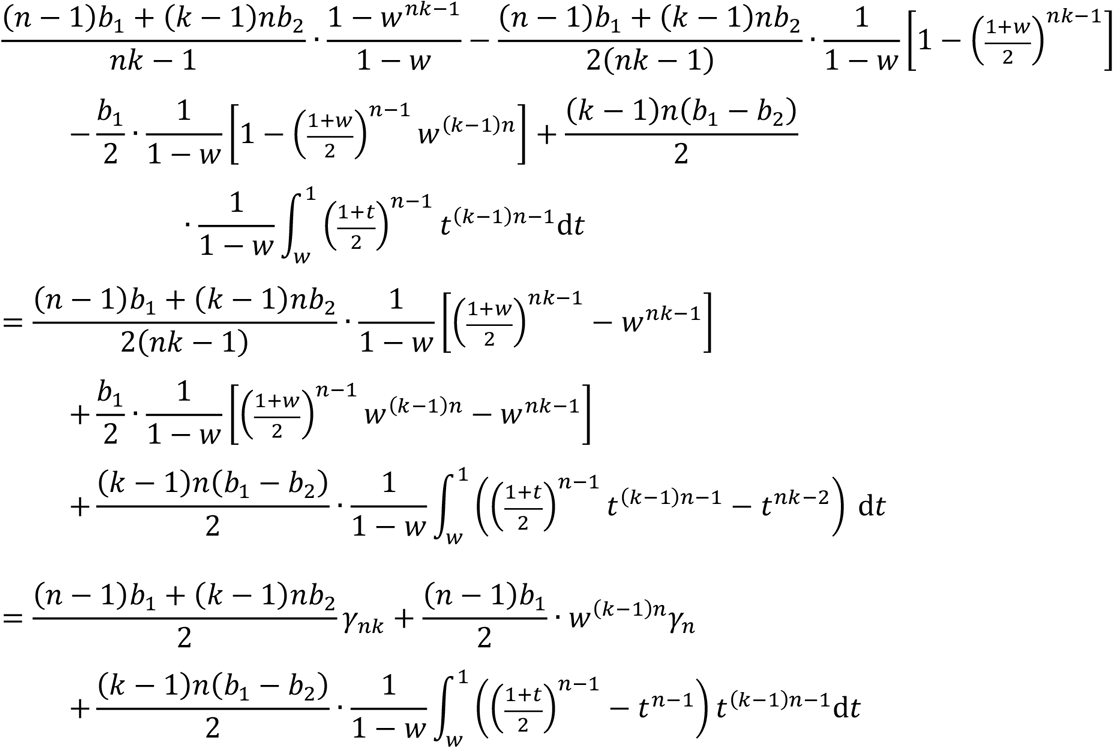

Introducing the notation

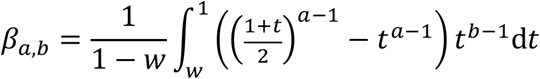

we arrive at the following characterization: Altruism is evolutionarily stable if

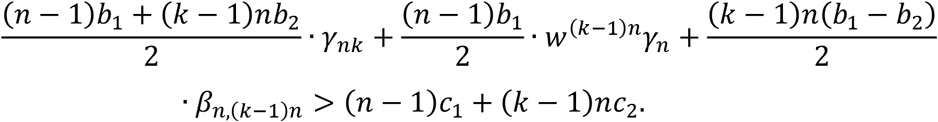

or equivalently,

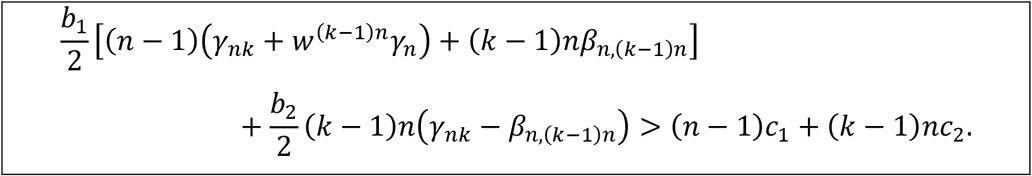

Note that 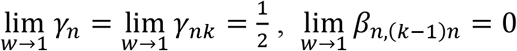. Thus, tending to 1 with *w*, we get back the condition of evolutionary stability in the additive model.

If there is no difference between the attitude toward full siblings or half-sibs, that is, *b*_1_ = *b*_2_ = *b* and *c*_1_ = *c*_2_ = *c*, then the condition for altruism to be evolutionarily stable is

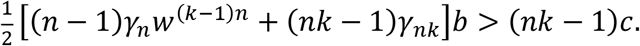

If altruism is bounded to full siblings only, that is, *b*_2_ = *c*_2_ = 0, then the use of the above *g* is no longer justifiable, because if half-sibs keep away from altruistic behavior, there is no point in keeping count of them in the group effect function *g*.

## Appendix 3. Haldane’s arithmetic and evolutionary stability

One can naturally ask whether there is any connection between our condition for evolutionary stability and Haldane’s arithmetic^7^.

Two possible cases can be distinguished:

a. Pairwise interactions. The effectiveness of rescue i.e., the survival probability *p* of an individual in a death trap, depends on each sibling’s willingness to help, but does not depend on the composition of the family. As we have already seen in Appendix 2, a special case of Haldane’s arithmetic, namely, Hamilton’s rule, is equivalent to the condition of evolutionary stability of altruism.
b. Group dependent survival rate. The increase of the survival probability *p* of an individual due to the contribution of each of her/his siblings is not fully determined by the sibling’s willingness to help: it also depends on the number of altruistic siblings. This dependence is quantified by the functions *f* and *g*.

Concentrating on the genes, let us adapt Haldane’s idea to our models. Since mutation can be assumed to be rare, pure altruistic families and mixed families (where one of the parents is heterozygote, carrying the selfish gene) determine the fitness of altruistic and selfish genes, resp., with but a negligible exception. That is, fitness is affected by the composition of the family, which is determined by the rules of Mendelian genetics.

Similarly to what we have done in Appendix 1, we are interested in the average number of altruistic genes rescued by an altruist when the composition of the group has an effect on the survival of those getting in danger.

Therefore, selecting a random parental gene, we ask whether the altruistic or the selfish variant will result in more surviving copies on the average, provided the resident population is homozygote altruist.^8^

Note that this is nothing else but what we have done in Appendix 2. There, evolutionary stability of altruism was defined by greater reproduction rate. The reproduction rate of the altruistic or selfish phenotype is just the average number of surviving offspring of the same type from one such parent selected uniformly at random. This can be obtained by summing the probabilities of being a same type survivor over all children, thus what we really have to do is to compare probabilities of inheriting the selected gene and surviving. To illustrate this, let us go through again the three mating models analyzed in Appendix 2. Note that every child inherits the fixed parental gene independently with probability 1/2, and this is also independent of the amount of support received from other siblings.

### Monogamous family

In this case, suppose a fixed parental gene is transferred to a child. If this gene is altruistic, the survival probability of the given child is increased by *bf*(*n* − 1) and decreased by *c*(*n* − 1). If the parental gene is selfish, then no cost decreases the survival probability, but the benefit from altruistic siblings is only *bf*(*ξ*), where *ξ* is a random variable having binomial distribution with parameters *n* − 1 and 1/2. Thus, it is worth being altruistic if *b*(*f*(*n* − 1) − 𝔼*f*(*ξ*)) > *c*(*n* − 1), just as in (1).

### Total promiscuity

In this case the selected parental gene can be maternal or paternal with equal probability. Suppose it is an altruistic gene. In the former case the increment of the survival probability of a fixed progeny is just the same as in the monogamous family, because (with overwhelming probability) all fathers are of resident type, and it is irrelevant if they are all different or a single person. In the latter case, i.e., when the focal gene is paternal, all siblings of the fixed child are from different fathers and they are altruistic. Thus, the change of the survival probability is negative: −*c*(*n* − 1). Taking expectation we get back the condition of (6),

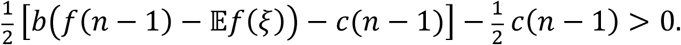

### Polygyny without promiscuity

Now, the selected parental gene is maternal or paternal with probabilities *k*/(*k* + 1) and 1/(*k* + 1), resp. When it is maternal, then a given child will have *N* = *n*_2_ + … + *n*_*k*_ altruistic half-sibs (without loss of generality we may suppose that *n* = *n*_1_ denotes the number of the focal mother’s children), and *n* − 1 altruistic full siblings in the altruistic case, to be compared with the selfish case where the number of altruistic full siblings is *ξ* again. Hence the increment of the survival probability is *g*(*n* − 1, *N*) − *c*(*n* − 1) vs. *g*(*ξ, N*). This affects *n*/2 children on average. This is also the case when the selected gene is paternal and happens to be altruistic, but it already concerns as many as *nk*/2 children on average. If we consider a heir of a selfish paternal gene, she/he will have a random number *η* of altruistic half-sibs; *η* is binomial with parameters *N* and 1/2, and, in addition, *ξ* altruistic full siblings. As there is no cost of altruism for her/him, the survival probability increases by *g*(*ξ, η*). Thus, altruism is remunerative if

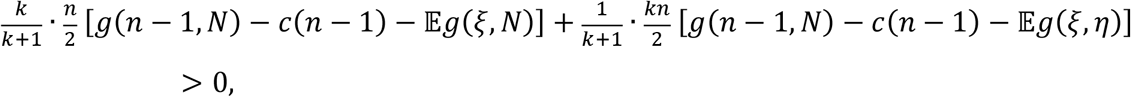

which is tantamount to (7).

Summing up, we found that Haldane’s arithmetic is equivalent to the condition of evolutionary stability in the presented population genetic models, for all different mating systems with pairwise interaction and group dependent survival rate, as well.

## Appendix 4. Generalized form of the classical Hamilton’s rule

As we have seen above, when there is no synergetic group effect, the classical Hamilton’s rule implies the evolutionary stability of sib altruism. However, in the presence of synergetic group effect, a generalized form of Hamilton’s rule is needed. To this end, let us take a look at our results from the viewpoint of Hamilton’s rule. Regarding the genetic relatedness in families, we have two cases:

### The genetic relatedness is uniform in the families

Under monogamy and total promiscuity, the genetic relatedness *r* between siblings is uniform, i.e. ½ and ¼, respectively. Observe that our conditions (3) and (6) can be read as *rbQ* > *c*, where

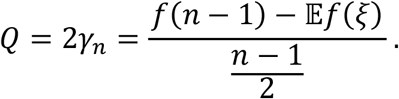

Thus, our generalized Hamilton’s rule takes account of the synergetic group effect with the multiplier *Q*, and this effect is different from the parameters of the classical Hamilton’s rule.

The numerator of *Q* is proportional to the difference between the average contribution of siblings’ help to the survival probability in a resident-resident family and a resident-mutant one. This difference is then normalized with the same quantity computed for the additive (*w* = 1) model, where no group effect takes place.

### The genetic relatedness is not uniform in the families

Under polygyny without promiscuity, each sib has full and half siblings, too. Moreover, each sib has perfect information whether his/her sib in danger is a full or a half sibling. Thus the altruist interaction can depend on the genetic relatedness of the interacting siblings. This makes our evolutionary stability condition most complicated (see Appendix 2.C.). Formally, under polygyny without promiscuity, our condition of the evolutionary stability of altruism reads as

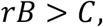

where

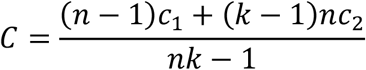

is the average cost; the average genetic relatedness within the families is

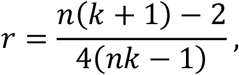

and the synergetic group effect combined with benefit parameters reads as

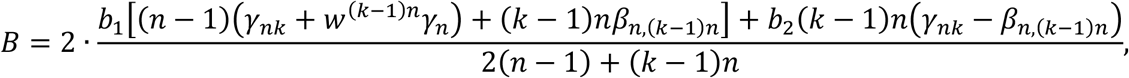

(see Appendix 2C.. for details). Here the discounted/synergetic group effect cannot be separated from the parameters of the classical Hamilton’s rule. In the additive model where the group does not take effect, 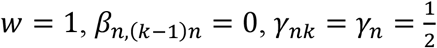, thus

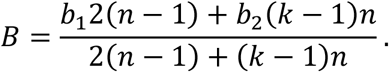

## Footnotes

1 The reader may doubt that the considered altruistic behaviour is really a moral rule. Here we follow the terminology of Bacha-Trams et al. (2017) who focused on the moral dilemma of Anna to donate one of her kidneys to her sister Kate, who is fatally ill with cancer. They found that the genetic relationship between interacting people robustly modulates social cognition of the perceiver in the human brain.

2 One may think that Haldane’s idea is only a piece of the biological folklore. However, Haldane (1955, on page 44) published his idea and he had the essential logic of Hamilton’s argument, see Maynard Smith (2017).

3 As we are looking for strict evolutionarily stable states, we do not consider neutral mutants.

4 Notice that if the actors are well-informed about the genetic relatedness between each other, then the interaction can depend on the genetic relatedness. Thus in general not only the benefit but also the cost can depend on *r*. In this case the possible Hamilton’s rule reads as *rB*(*r*) > *C*(*r*).

5 Observe that this condition is just Hamilton’s rule.

6 Thus in pairwise interactions Hamilton’s rule is valid again.

7 Note that the individuals are not in a decision-making situation, as their behavior is completely determined by their genes.

8 Consider a mutant gene. There are two types of mutation: the new mutant gene is also altruistic, but its genetic relatedness with the resident is 0, or the mutant gene is not altruistic.

